## Supplementary Information for "Real-time 3D Single Molecule Tracking"

| **Section** | **Page** |
| --- | --- |
| **Figures** |  |
| Supplementary Figure 1: Schematic of 3D-SMART setup | 5 |
| Supplementary Figure 2: Piezo stage response | 6 |
| Supplementary Figure 3: Tracking simulation | 7 |
| Supplementary Figure 4: Optimization of tracking parameters via simulation | 8 |
| Supplementary Figure 5: Schematic of photon antibunching measurement | 9 |
| Supplementary Figure 6: Trajectories of molecules used in antibunching measurement | 10 |
| Supplementary Figure 7: Additional antibunching data of single Atto 647N dye in 90% glycerol. | 11 |
| Supplementary Figure 8: Tracking "on-time" analysis | 12 |
| Supplementary Figure 9: Photons per molecule with tracking disabled | 13 |
| Supplementary Figure 10: BSA-Atto 647N photo-blinking | 14 |
| Supplementary Figure 11: Single Alexa 488 tracking in 90 wt% glycerol | 15 |
| Supplementary Figure 12: BSA-Alexa 488 tracking in 73.5 wt% glycerol | 16 |
| Supplementary Figure 13: Characterization of DNA substrates | 17 |
| Supplementary Figure 14: Tracking a range of dsDNA lengths | 19 |
| Supplementary Figure 15: Distribution of diffusion coefficients of various lengths of dsDNA. | 20 |
| Supplementary Figure 16: Comparison of measured diffusion coefficients of various lengths of dsDNA to theoretical models | 22 |
| Supplementary Figure 17: Histograms of trajectory durations | 24 |
| Supplementary Figure 18: Histogram of diffusion coefficients of mRNA-DFHBI-1T | 26 |
| Supplementary Figure 19: Ensemble probe ssDNA binding to mRNA | 27 |
| Supplementary Figure 20: Alexa 488 bleaching time | 28 |
| Supplementary Figure 21: fastFISH probe DNA non-specific binding to dsDNA | 29 |
| Supplementary Figure 22: 2D scatter plot of photon counts of mRNA channel versus diffusion coefficient of template double strands DNA | 30 |
| Supplementary Figure 23: Sample holder used for 3D-SMART | 31 |
| Supplementary Figure 24. Tracking precision. | 32 |
| Supplementary Figure 25. Scatter plot of diffusion coefficient and durations | 33 |
| **Tables** |  |
| Supplementary Table 1: Comparison of RT-3D-SMT methods | 34 |
| Supplementary Table 2: Comparison of trajectory duration and photons per molecule with active feedback on and off | 35 |
| **Notes** |  |
| Supplementary note 1: Antibunching measurement | 36 |
| Supplementary note 2: Diffusion coefficient calculation | 37 |
| Supplementary note 3: 2D kernel density estimation | 38 |
| Supplementary note 4: Stage calibration | 39 |
| Supplementary note 5: Factors will affect collected signal intensity | 40 |
| **Movie descriptions** | 41 |
| **Supplementary Codes: Tracking simulation** | 42 |
| **Supplementary data** | 47 |

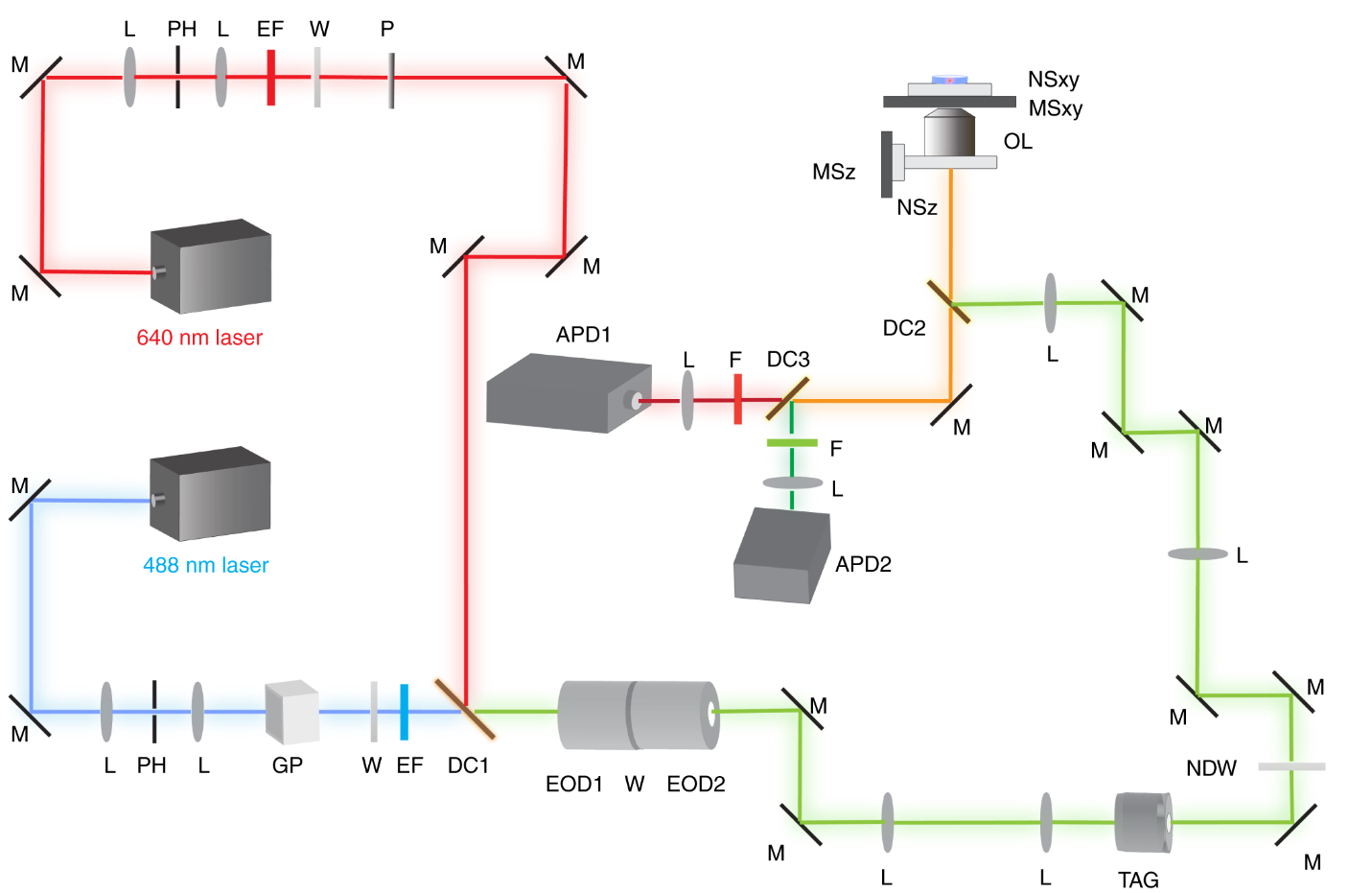

**Supplementary Figure 1.** Schematic of 3D-SMART setup. L: lens; M: mirror; PH: pinhole; GP: Glan-Thompson polarizer; P: polarizer; W: half waveplate; EF: excitation filter; DC: dichroic mirror; EOD: electro-optic deflector; TAG: TAG lens; NDW: Neutral density filter wheel; OL: objective lens; MSxy: xy microstage; MSz: z microstage; NSxy: xy nanopositioner stage; NSz: z nanopositioner stage; F: fluorescence filter.

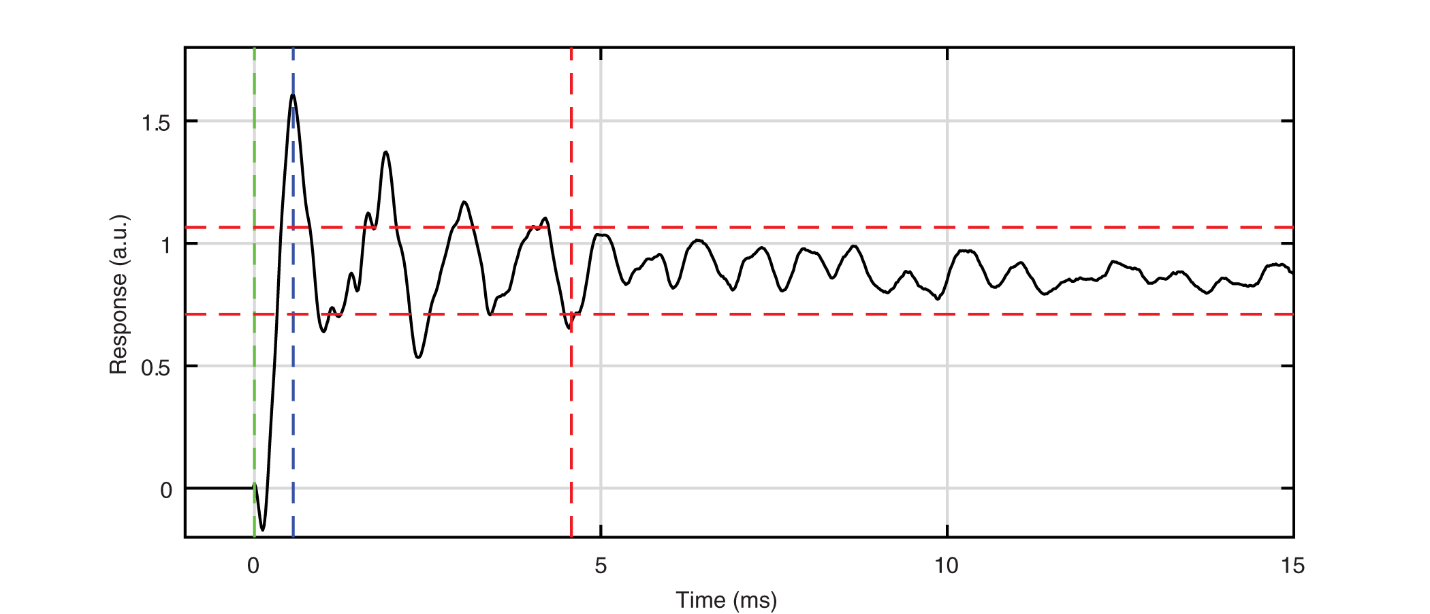

**Supplementary Figure 2**. Piezo stage (MadCity Labs Nano-LPQ) response to a step of a 1 micron. Step occurs at t = 0 ms (green dotted line). Peak time is 0.56 ms (blue dotted line) and the 20% settling time is 4.57 ms (red dotted line).

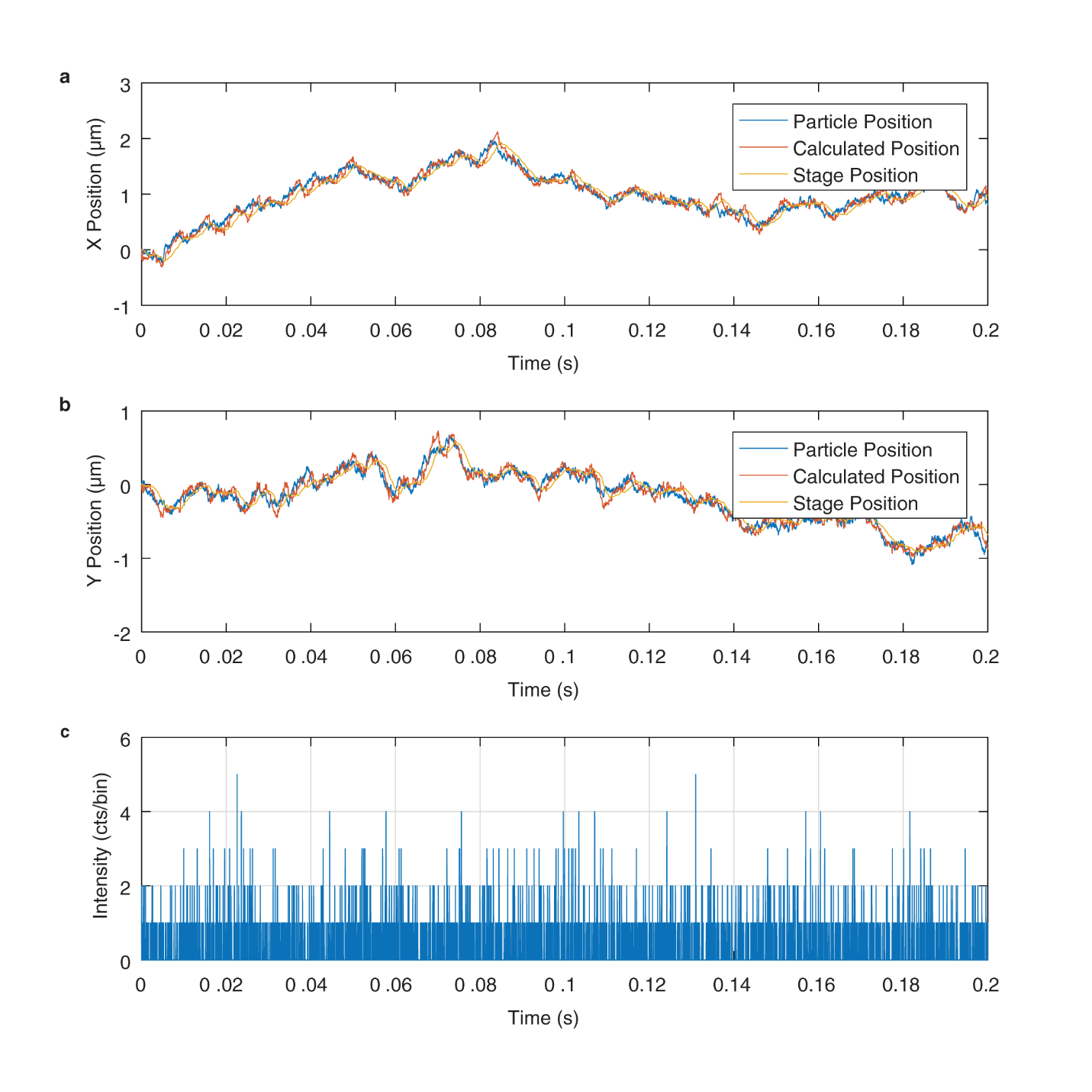

**Supplementary Figure 3.** Tracking Response simulation. Simulated 2D Tracking Response in (**a**) X and (**b**) Y using the ‘model1Track’ code provided below. (**c**) Intensity from simulated trajectory.

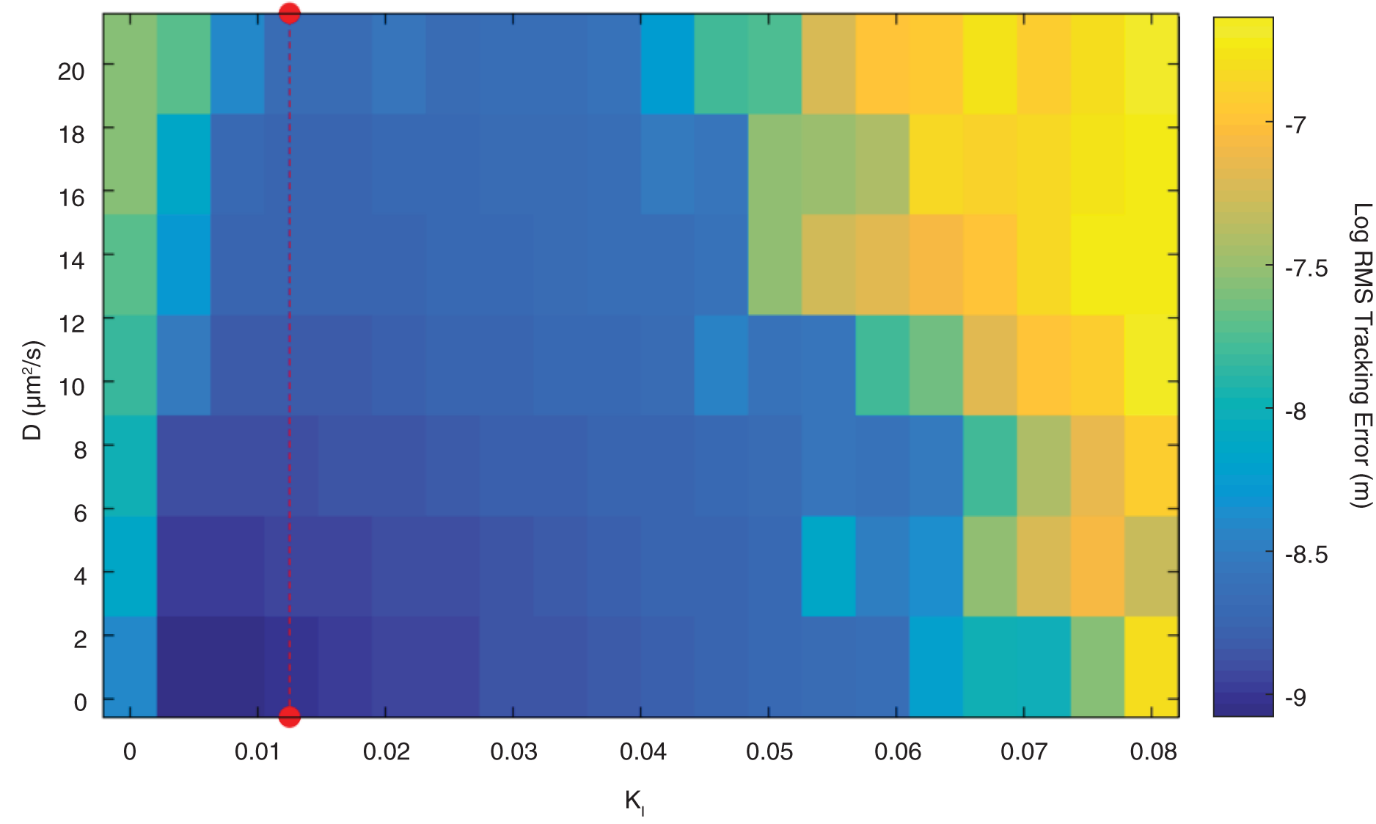

**Supplementary Figure 4.** Optimization of Tracking Parameters via Simulation. To predict the best feedback parameters for RT-3D-SMT, simulated 2D tracking was performed for a variety of diffusion coefficients (D) and integral feedback values (K_I_) using the model1Track code shown below. The average count rate from a simulated tracked molecule was ~ 10 kHz and a background count rate of ~ 700 Hz. Data are evaluated based on the RMS tracking error, the difference between the actual particle position and the piezo stage coordinates. The red dot line indicates the best K_I_ predicted for SMT, a value of ~ 0.012, displaying excellent tracking performance across a wide range of diffusion coefficients.

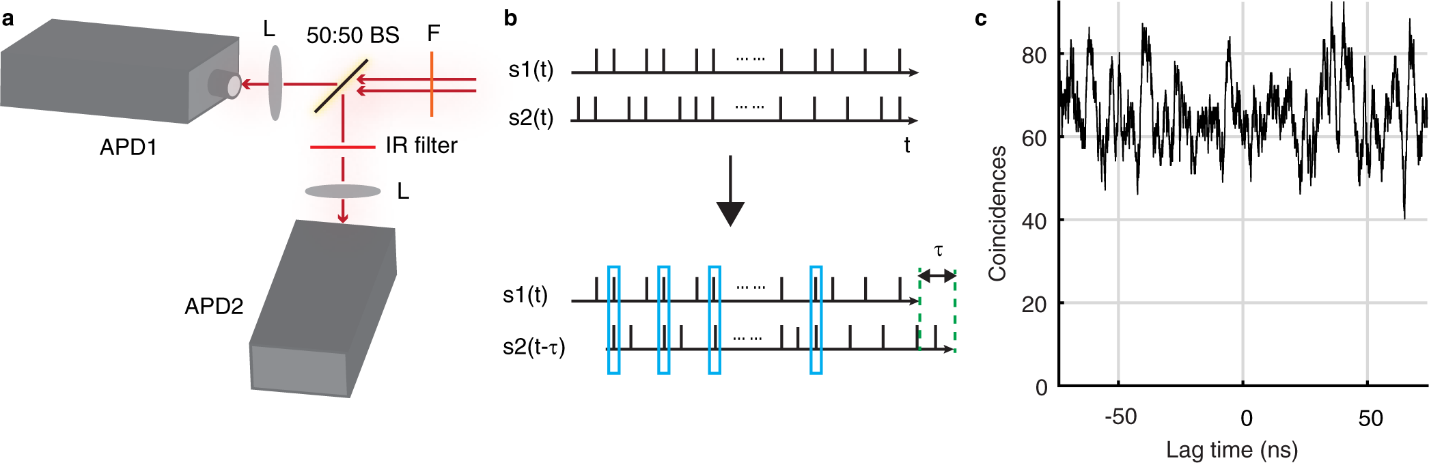

**Supplementary Figure 5.** Schematic of photon antibunching measurement. (**a**) Schematic of fluorescence detection by two APDs. (**b**) Photon pair correlation calculation. The photons arrival times of each APD are tagged by TCSPC. The Photon pair correlation is calculated as the sum of coincidence after shifting one signal with lag time τ. (**c**) Photon pair correlation coincidence for a fluorescent bead tracking (bin time: 2 ns).

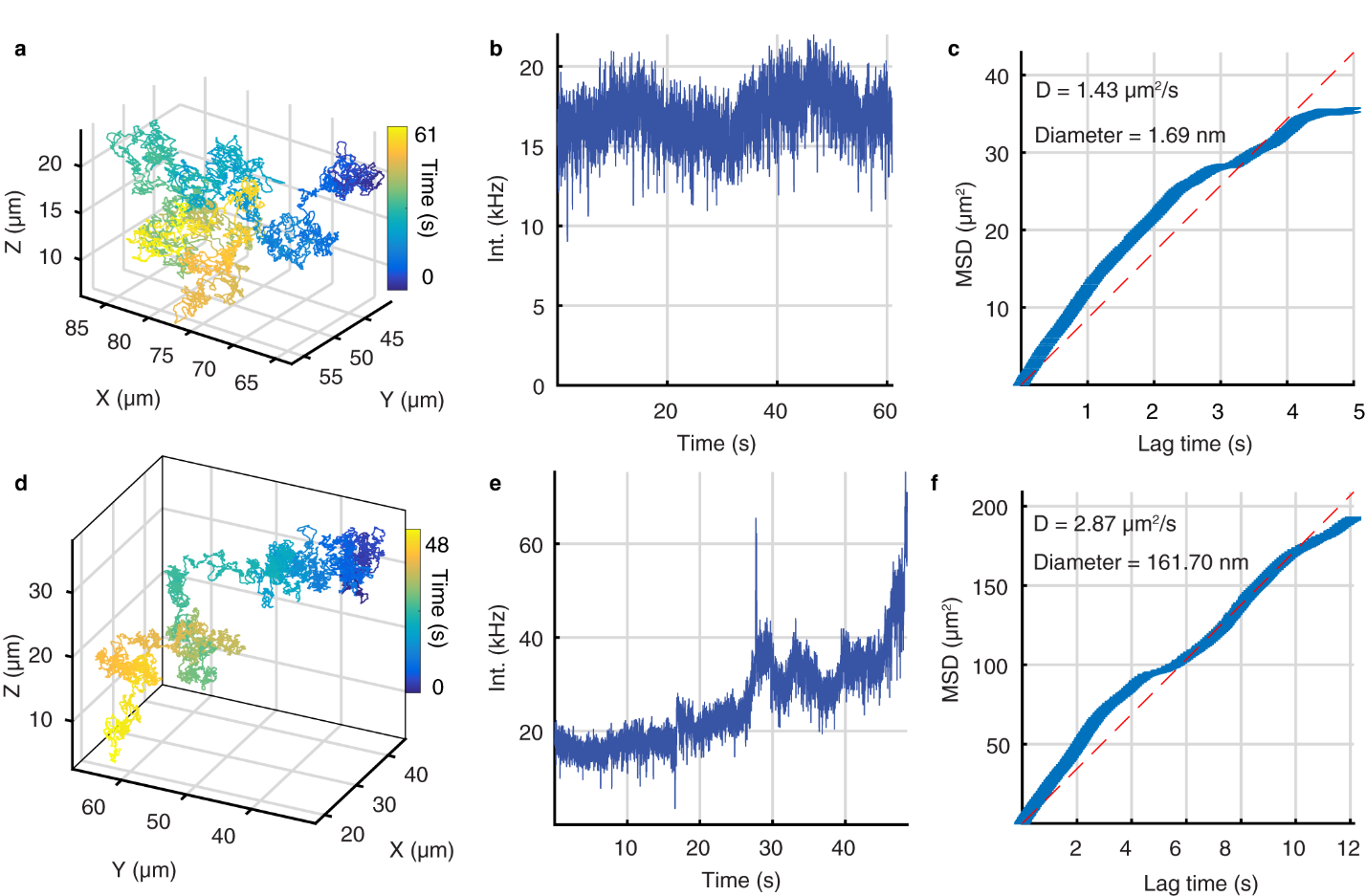

**Supplementary Figure 6.** Trajectories used in antibunching measurement. (**a**-**c**) Trajectory, intensity and mean square displacement of **Fig. 2e**. (**d**-**f**) Trajectory, intensity and mean square displacement of **Supplementary Fig. 5c**.

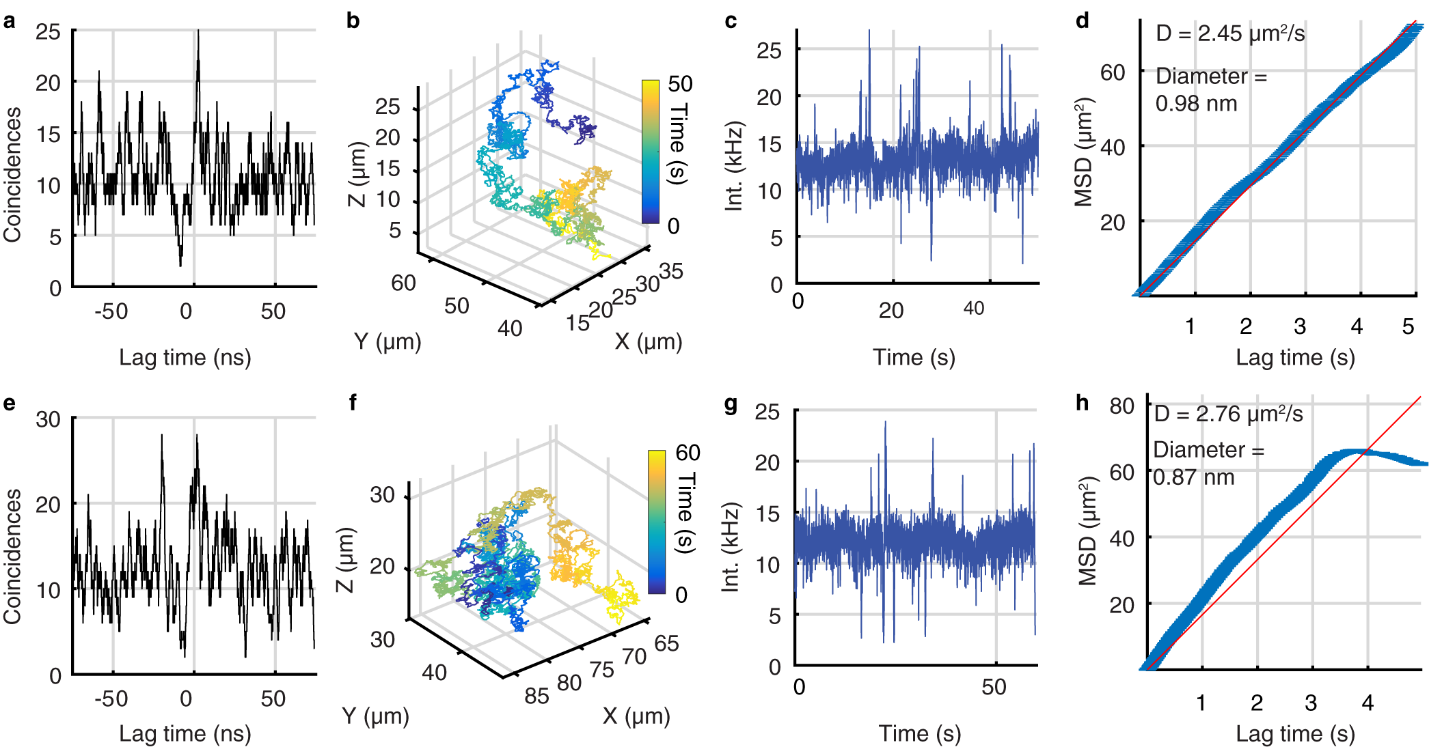

**Supplementary Figure 7.** Additional antibunching data of single Atto 647N dye in 90% glycerol. (**a**) Photon pair correlation coincidence for trajectory (**b**). Bin time: 2 ns. (**c**, **d**) Intensity trace and mean square displacement of trajectory (**b**). (**e**) Photon pair correlation coincidence for trajectory (**f**). (**g**, **h**) Intensity trace and mean square displacement of trajectory (**f**).

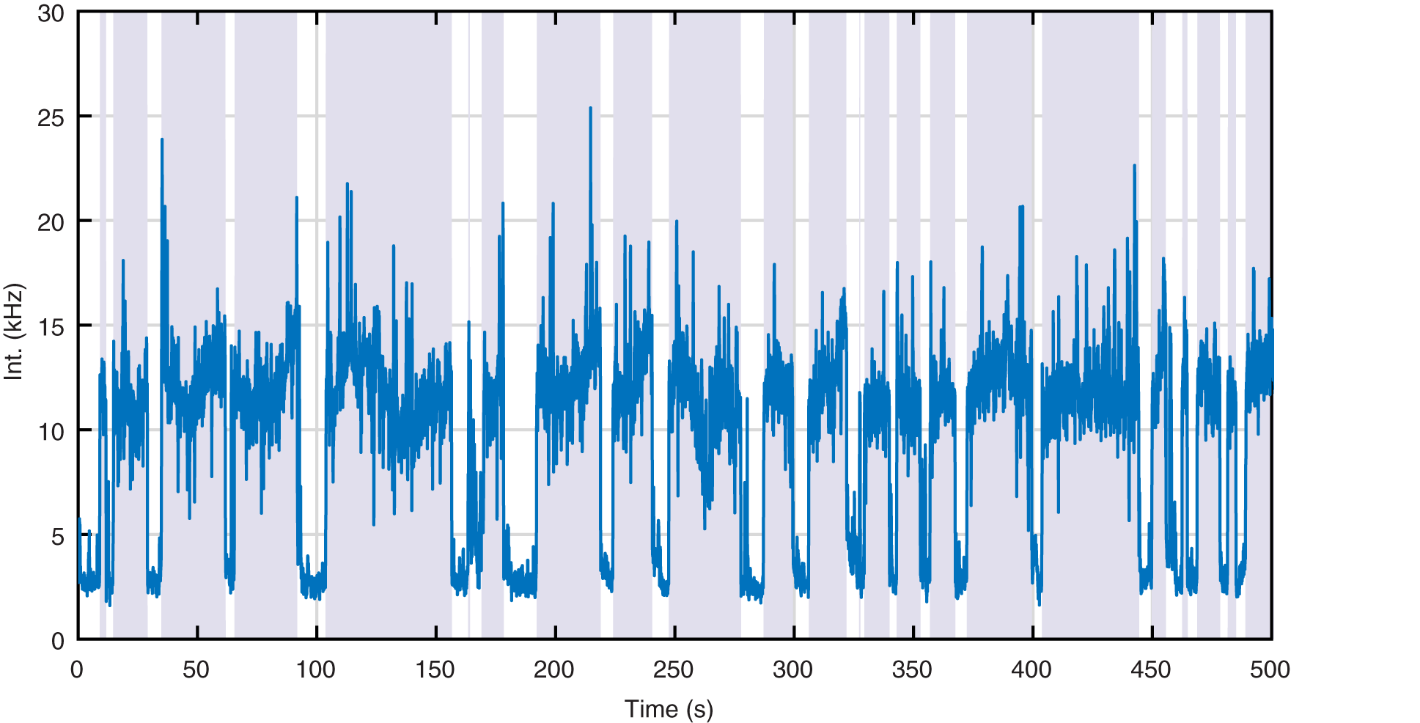

**Supplementary Figure 8.** Tracking "on-time" analysis. The gray bars represent when the tracking mechanism is actively tracking a molecule. Over this 500 s period, the tracking system is collecting data 72.9% of the time, with a mean trajectory length of 16.0 ± 13.4 s. These data can also be seen in **Supplementary Movie 2**.

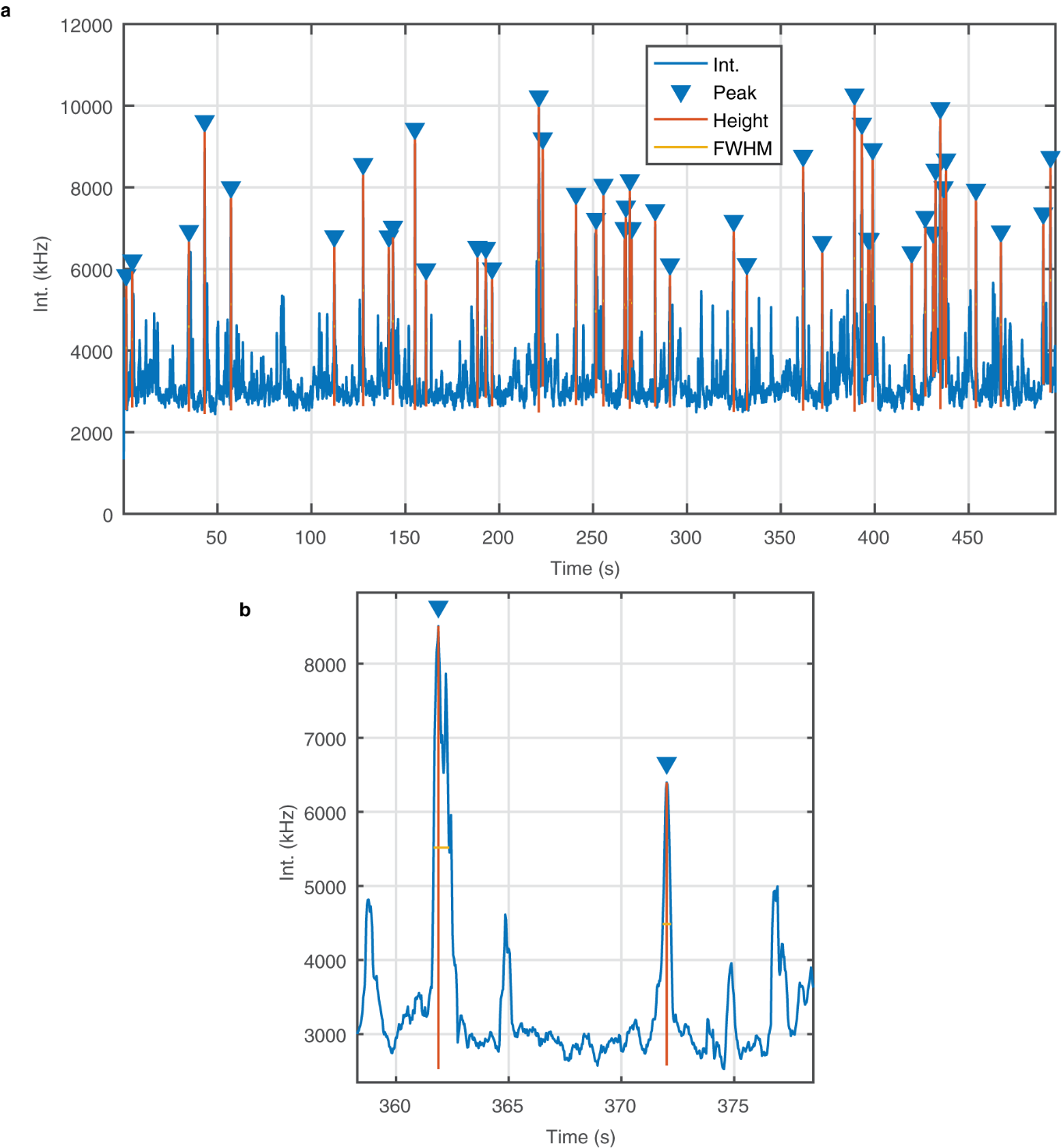

**Supplementary Figure 9.** Photons per molecule with tracking disabled. The MATLAB function findpeaks was used to find the peak height and FWHM for peaks exceeding 5 kHz. The average FWHM was 0.457 ± 0.169 s.

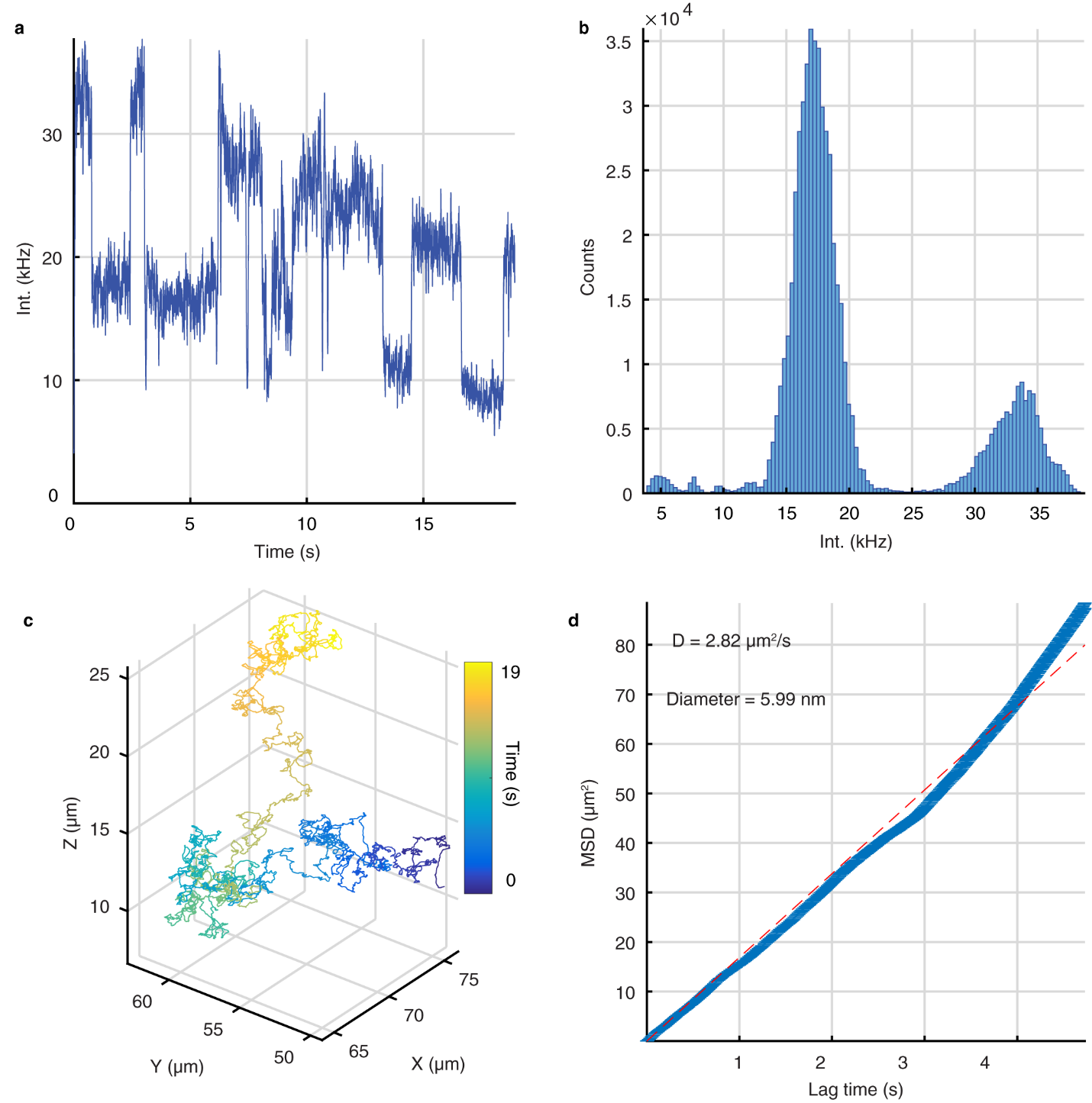

**Supplementary Figure 10.** BSA-Atto 647N photo-blinking whilst tracking. (**a**) Fluorescent intensity as a function of time for the trajectory in (**c**). (**b**) Intensity distribution of first 6 s of the trajectory shows two peaks, the higher intensity peak corresponding to both dyes in an emissive state while the lower intensity peak corresponds to one of the dyes in a dark state. (**d**) MSD of the trajectory in (**c**).

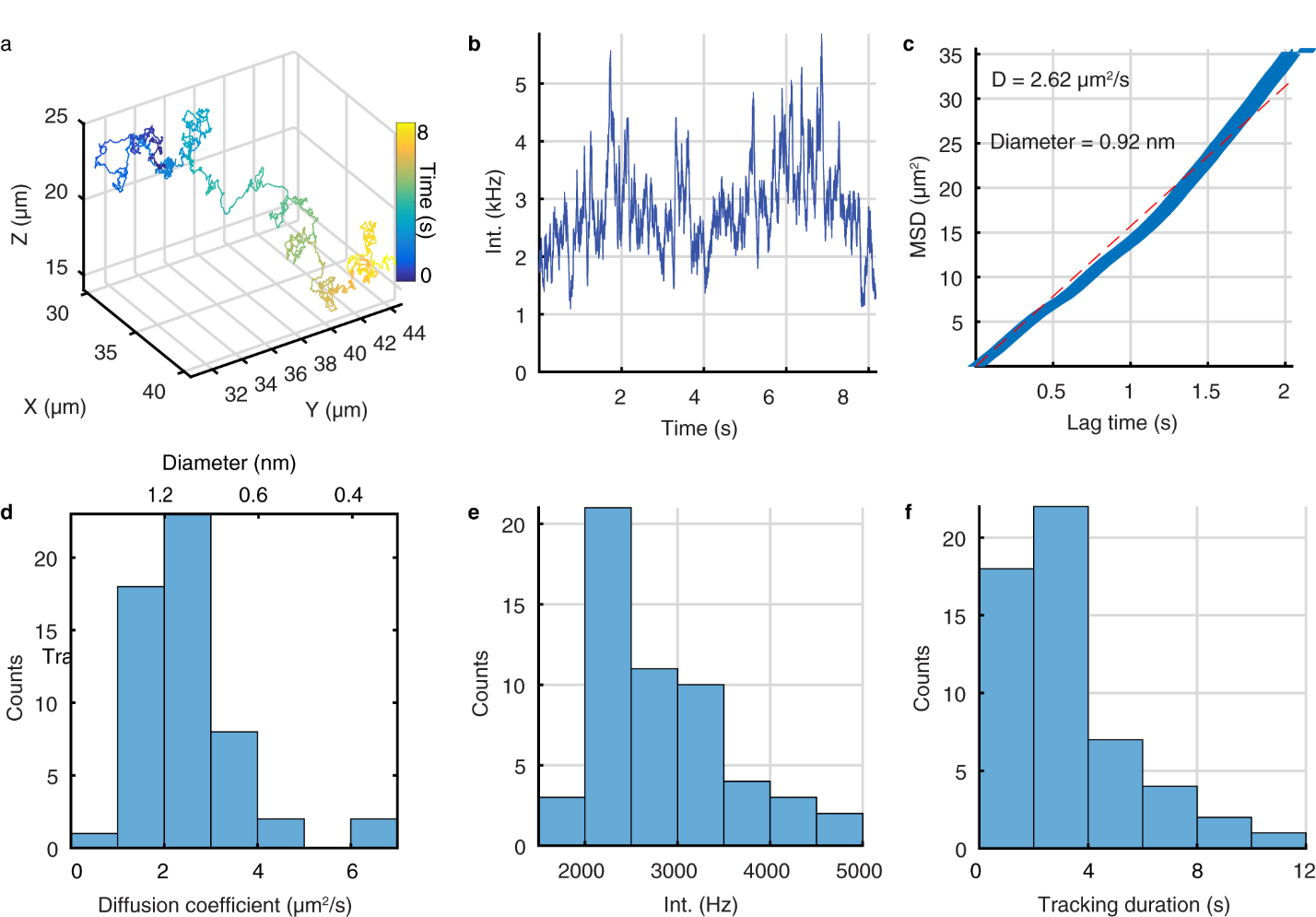

**Supplementary Figure 11.** Single Alexa 488 tracking in 90 wt% glycerol. (**a**) 3D trajectory of a freely diffusing single Alexa 488 molecule 90% glycerol solution. (**b**) Fluorescent intensity as a function of time for the trajectory in (**a**). (**c**) MSD of the trajectory in (**a**). The blue line is the measured MSD while the dotted red line is best fit line from linear regression. (**d**-**f**) MSD (**d**), mean intensity and tracking length (**f**) analysis of 54 trajectories. The mean intensity was measured over the first 100 ms of each trajectory. The mean tracking duration is 3.25 ± 2.18 s.

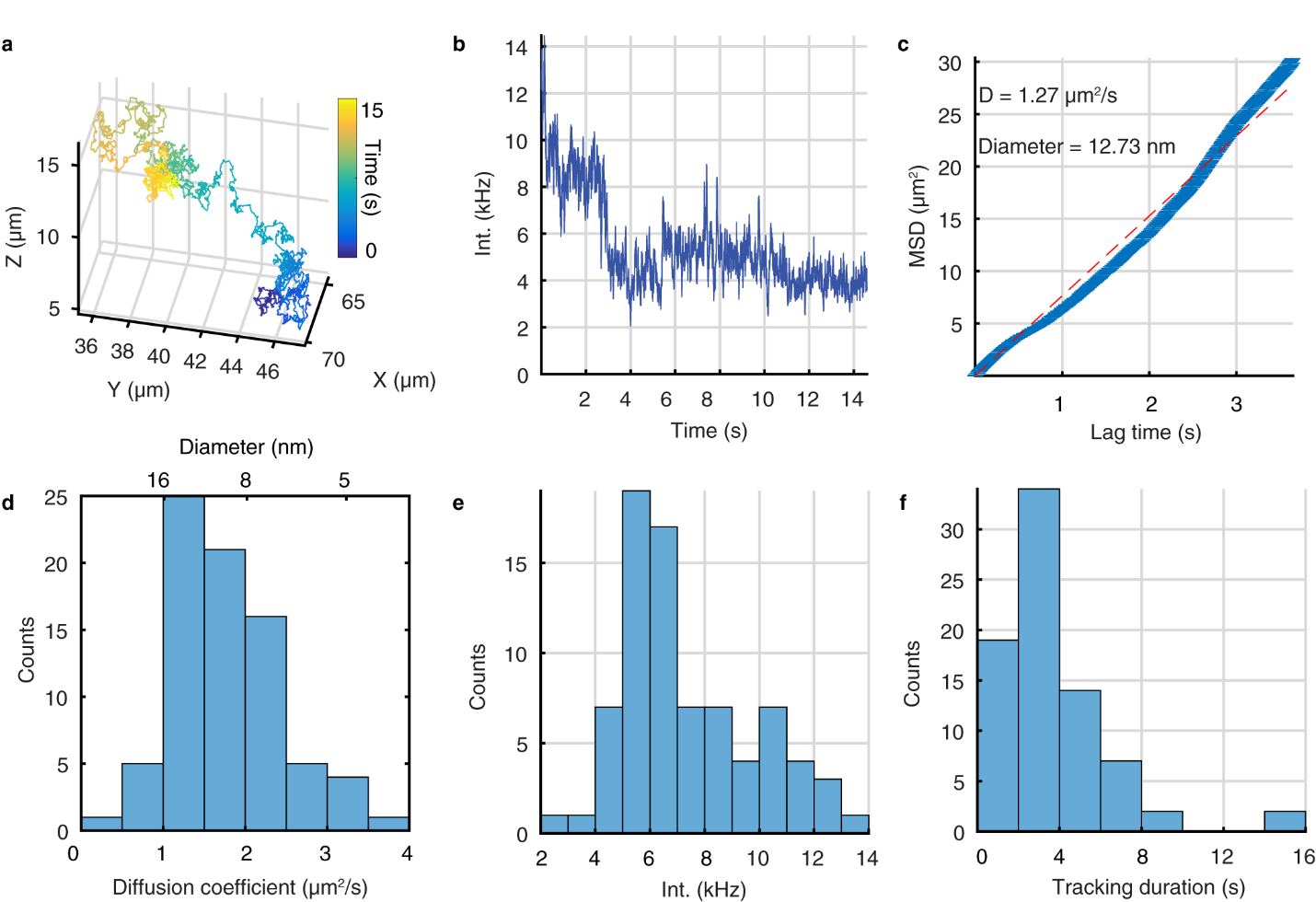

**Supplementary Figure 12.** BSA-Alexa 488 tracking in 73.5 wt% glycerol. (**a**) 3D trajectory of a freely diffusing single BSA molecule 73.5 wt% glycerol solution. (**b**) Fluorescent intensity as a function of time for the trajectory in (**a**). (**c**) MSD of the trajectory in (**a**). The blue line is the measured MSD while the dotted red line is best fit line from linear regression. (**d**-**f**) MSD (**d**), mean intensity (**e**) and tracking duration (**f**) analysis of 78 trajectories. The mean intensity was measured over the first 100 ms of each trajectory. The mean tracking duration is 3.78 ± 2.67 s.

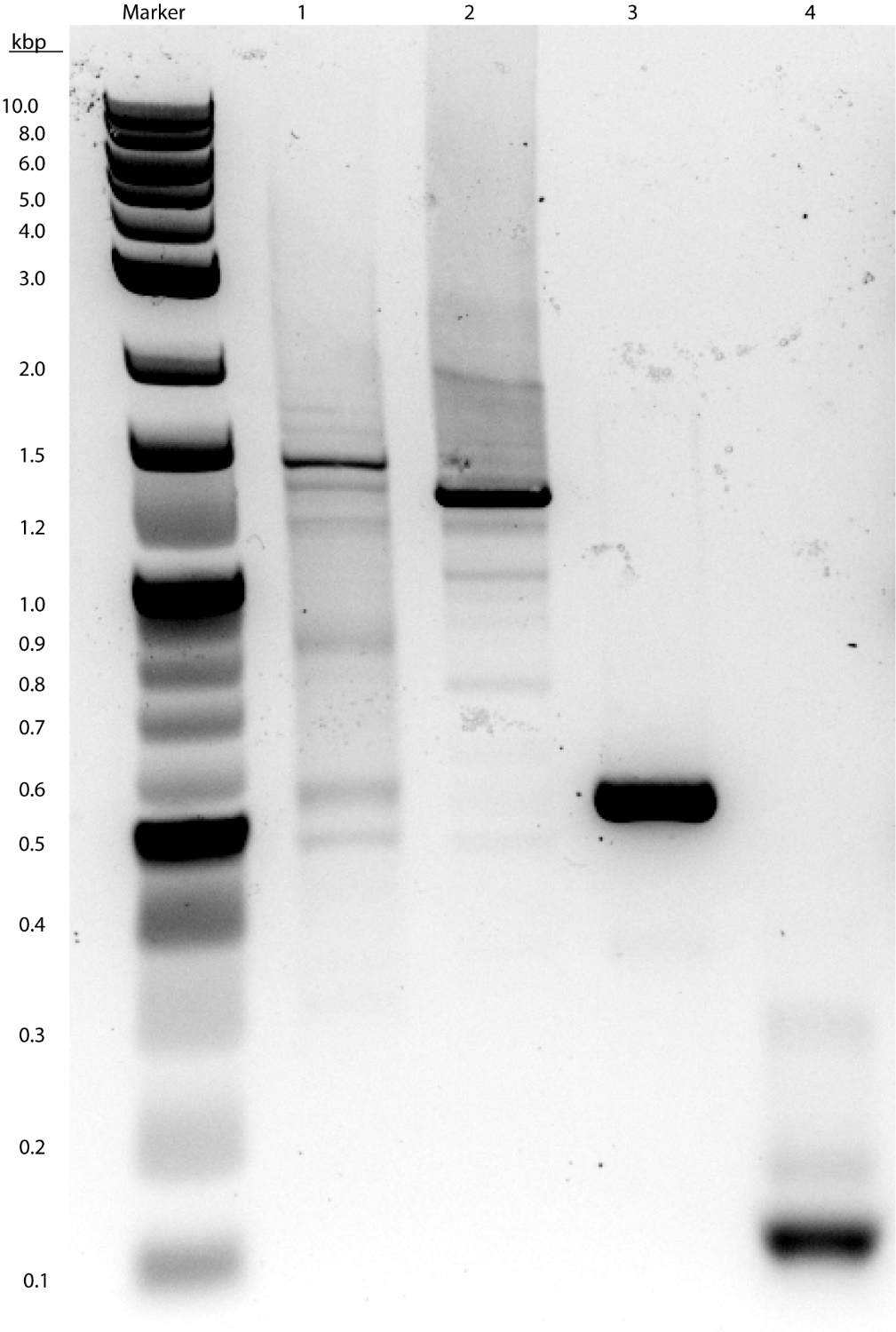

**Supplementary Figure 13.** Characterization of DNA substrates used in single molecule tracking and transcription experiments. Analysis of Atto 647N-labeled DNA substrates performed by electrophoresis. 1385 bp, 1218bp, and 547 bp dsDNA produced by PCR (lanes 1-3), and hybridization of synthesized oligonucleotides to produce 99 bp dsDNA (lane 4). DNA samples (200 ng) were separated on 1.5 % Omnipur agarose gel (EMD), with 1X GelRed (Biotium) nucleic acid stain, in 1X TAE, pH 8.3 at 8 V/cm and directly imaged on Chemidoc Imager (BioRad) at 302 nm. Marker, 1 kb Plus DNA Ladder (NEB).

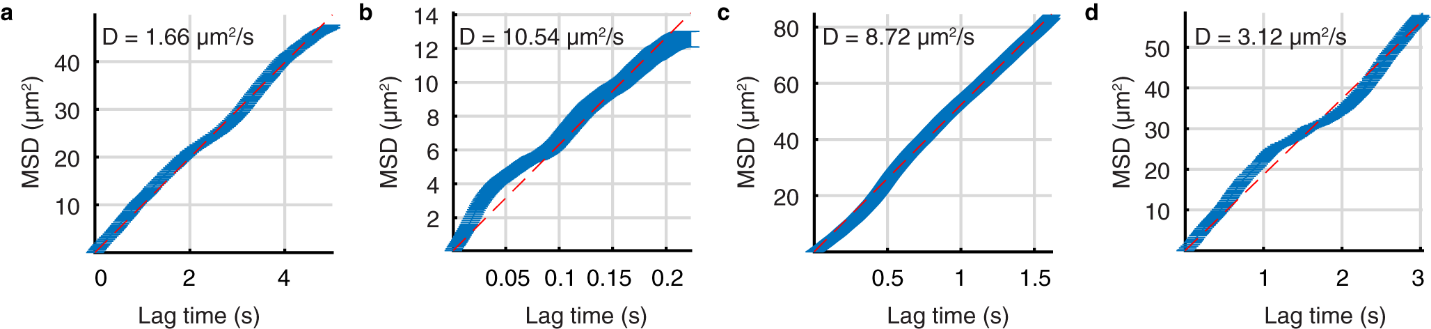

**Supplementary Figure 14.** Tracking a range of dsDNA lengths at indicated glycerol concentrations. (**a**) MSD of the trajectory in **Fig. 2d**. (**b**) MSD of the trajectory in **Fig. 2e**. (**c**) MSD of the trajectory in **Fig. 2f**. (**d**) MSD of the trajectory in **Fig. 2g**. The blue line is the measured MSD while the dotted red line is best fit line from linear regression.

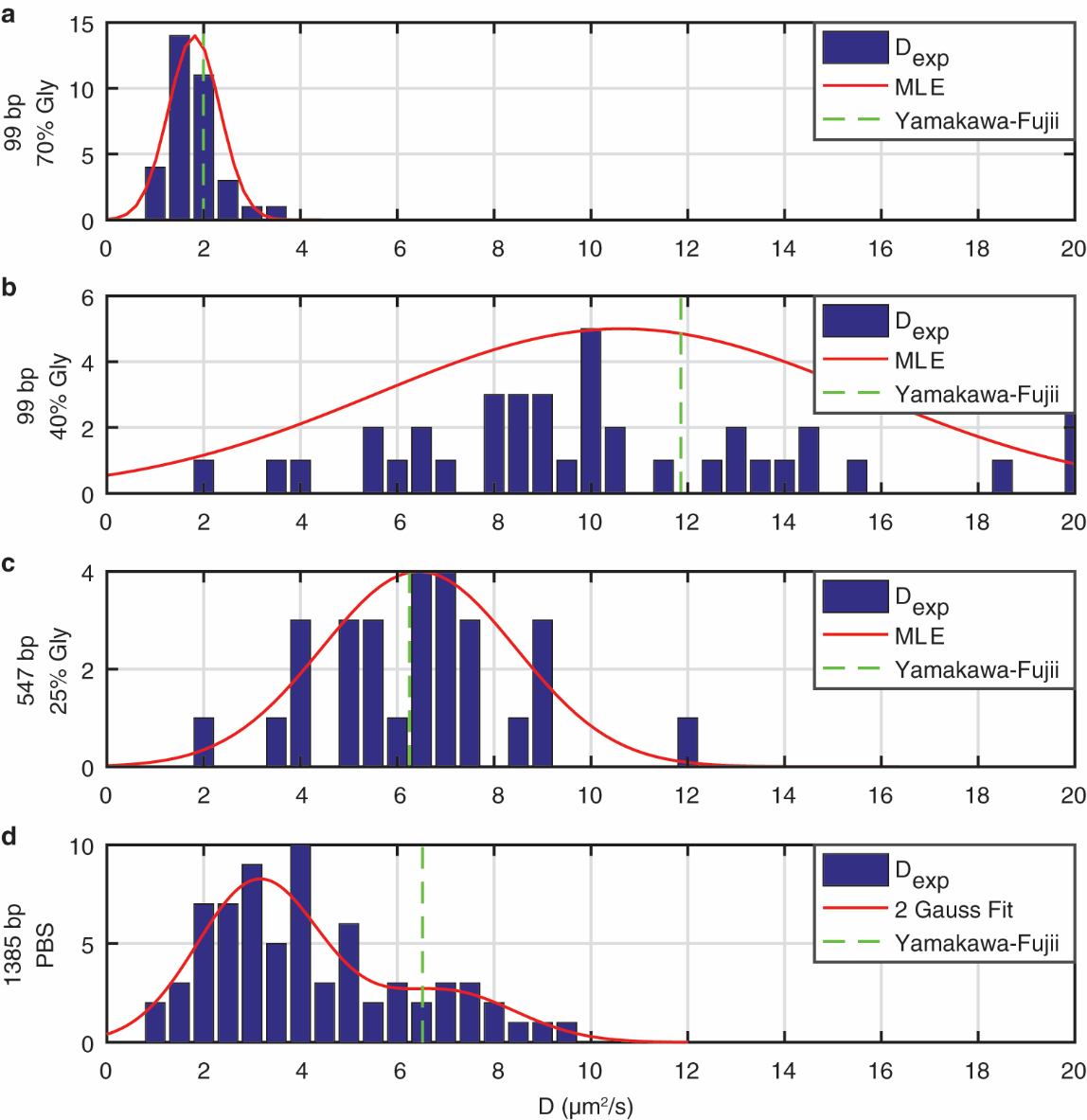

**Supplementary Figure 15.** Distribution of diffusion coefficients of various lengths of dsDNA. Diffusion coefficient distribution for (**a**) 99 bp dsDNA tracked in 70 wt% glycerol, (**b**) 99 bp dsDNA tracked in 40 wt% glycerol, and (**c**) 547 bp dsDNA tracked in 25 wt% glycerol. The red curves show the best fit Gaussian distribution calculated via the maximum likelihood method. The green dotted line represents the expected diffusion coefficient from a wormlike chain model with a persistence length of 50 nm and a diameter of 2.5 nm^37^. (**d**) 1385 bp dsDNA tracked in PBS. The red curve shows the best fit of the distribution to a sum of two Gaussians. The green dotted line represents the expected diffusion coefficient from the wormlike chain model developed by Yamakawa and Fujii^37^. It should be noted that the wider distribution of the diffusion coefficient for the 99bp dsDNA at 40 wt% compared to 70 wt% glycerol is due to the uncertainty in the measured diffusion coefficient at higher diffusive speeds, not a change in the molecular conformation."

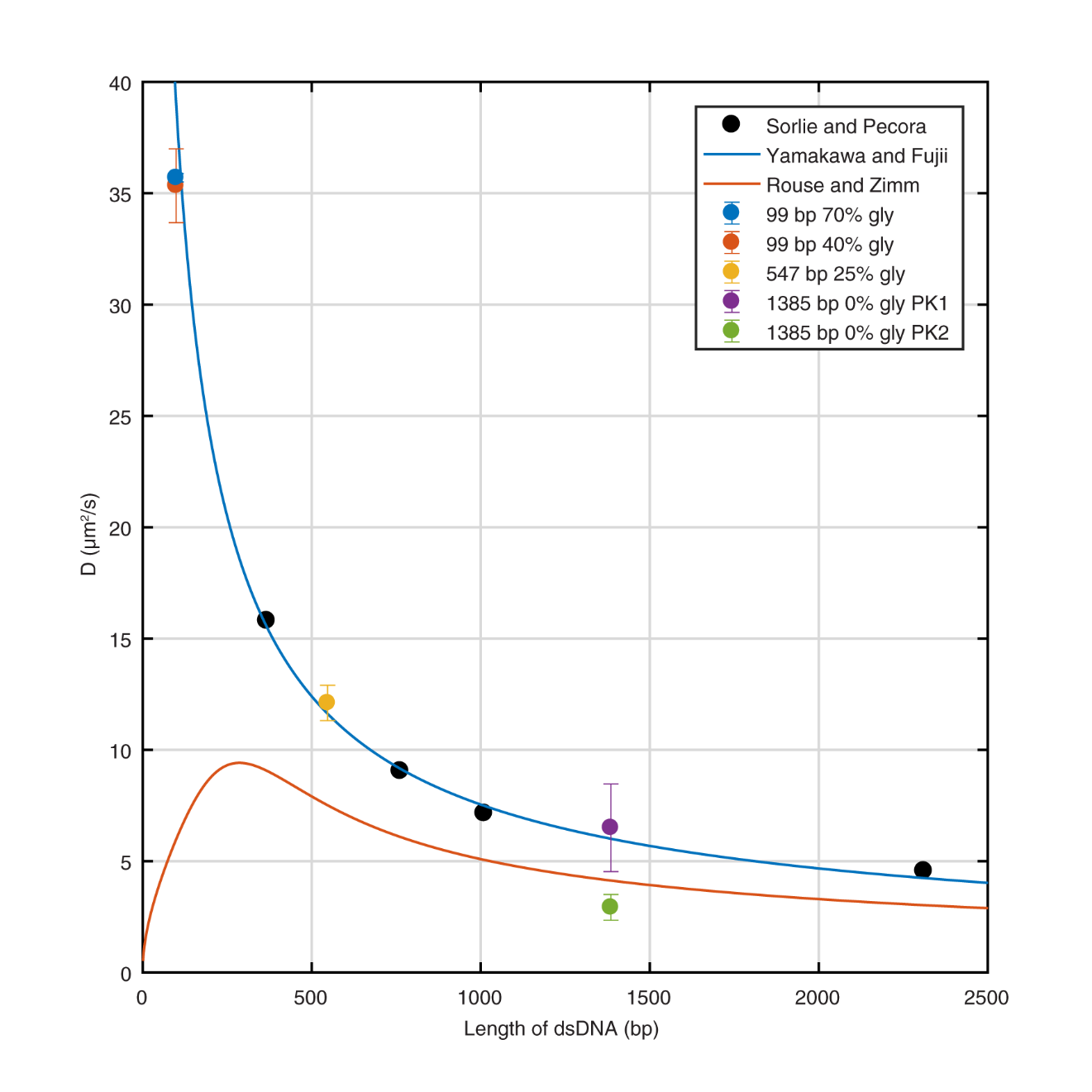

**Supplementary Figure 16.** Comparison of measured diffusion coefficients from active 3D tracking to theoretical models and previous experimental data. To facilitate comparison, the diffusion coefficients measured by active feedback 3D tracking are scaled to the diffusive behavior in water by multiplying by the ratio of the viscosity of the experimental glycerol solution to that of the viscosity of water ($\eta_{gly}/\eta_{water})$. For the 1385 bp sample, the two different peaks extracted from the distribution are plotted individually (PK1 and PK2). For comparison, the orange curve shows the expected diffusion coefficient predicted by the Rouse and Zimm model with the radius of gyration calculated using a wormlike chain model^38^. The blue curve shows the wormlike chain treatment using the Oseen-Burgers procedure with a persistence length of 50 nm and a diameter of 2.5 nm^37^. For comparison with light scattering data, the black circles show the experimental results from Sorlie and Pecora on DNA restriction fragments of varying lengths^38^. Note that the active 3D tracking results are in excellent agreement with the wormlike chain model of Yamakawa and Fujii as well as the experimental data of Sorlie and Pecora across a wide range of dsDNA lengths.

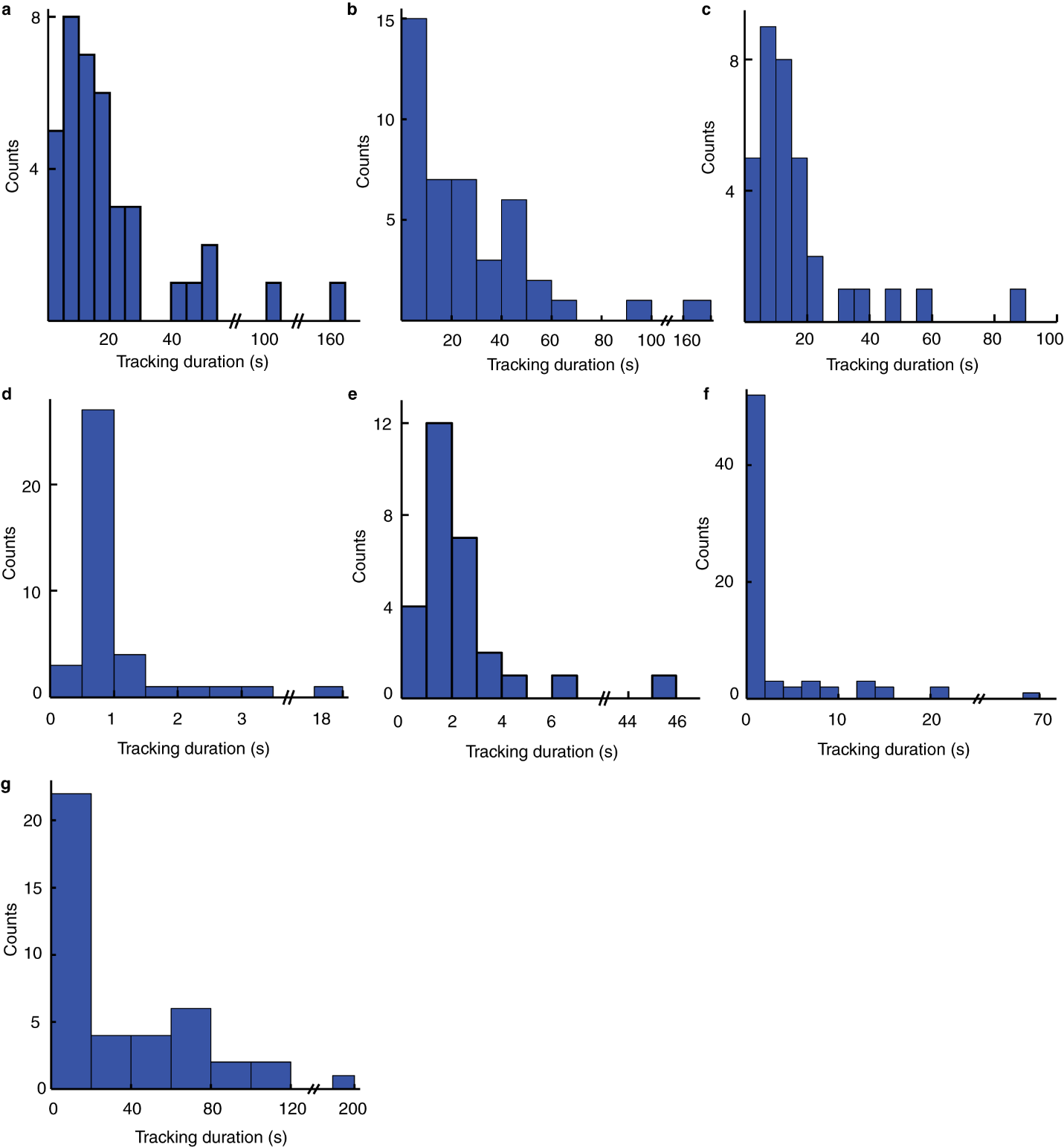

**Supplementary Figure 17.** Histograms of tracking trajectory duration times. Histograms of tracking trajectory duration times for single Atto 647N NHS ester in 90 wt% glycerol-water solution (**a**), Atto 647N labeled BSA in 73.5 wt% glycerol-PBS solution (**b**), 99 bp dsDNA tracking in 70 wt% glycerol solution (**c**), 99 bp dsDNA tracking in 40 wt% glycerol solution (**d**), and 547 bp dsDNA tracking in 25 wt% glycerol solution (**e**), single Atto 647N labeled dsDNA in PBS solution (**f**), and mRNA-DFHBI-1T in 75 wt% glycerol-binding buffer solution (**g**). The mean tracking duration is 22.69± 29.87 s (**a**), 30.126 ± 41.34 s (**b**), 17.11 ± 17.60 s (**c**), 1.34 ± 2.78 s (**d**), 3.71 ± 8.32 s (**e**), 3.88 ± 9.29 s (**f**) and 36.74 ± 41.51 s (**g**).

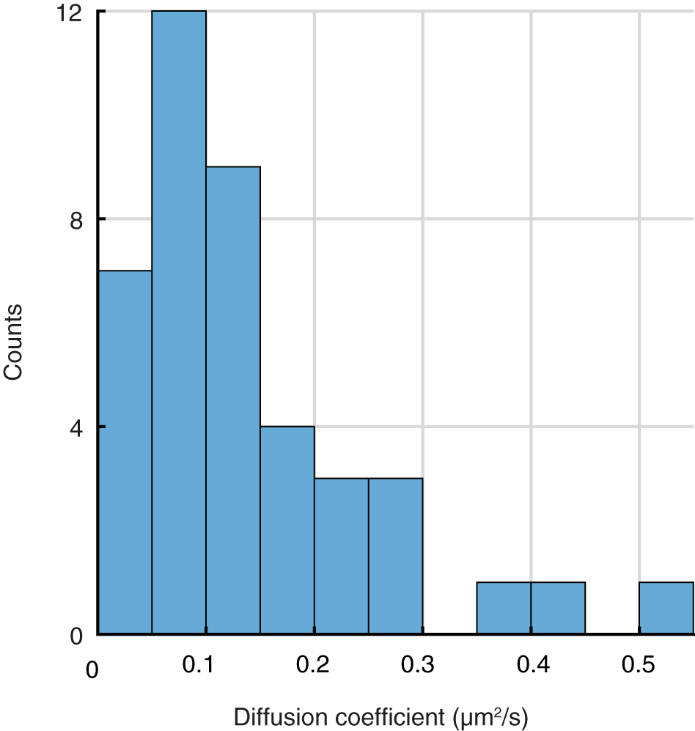

**Supplementary Figure 18.** Histogram of diffusion coefficient of mRNA-DFHBI-1T. The mean diffusion coefficient is 0.14 ± 0.11 µm^2^/s. n = 41.

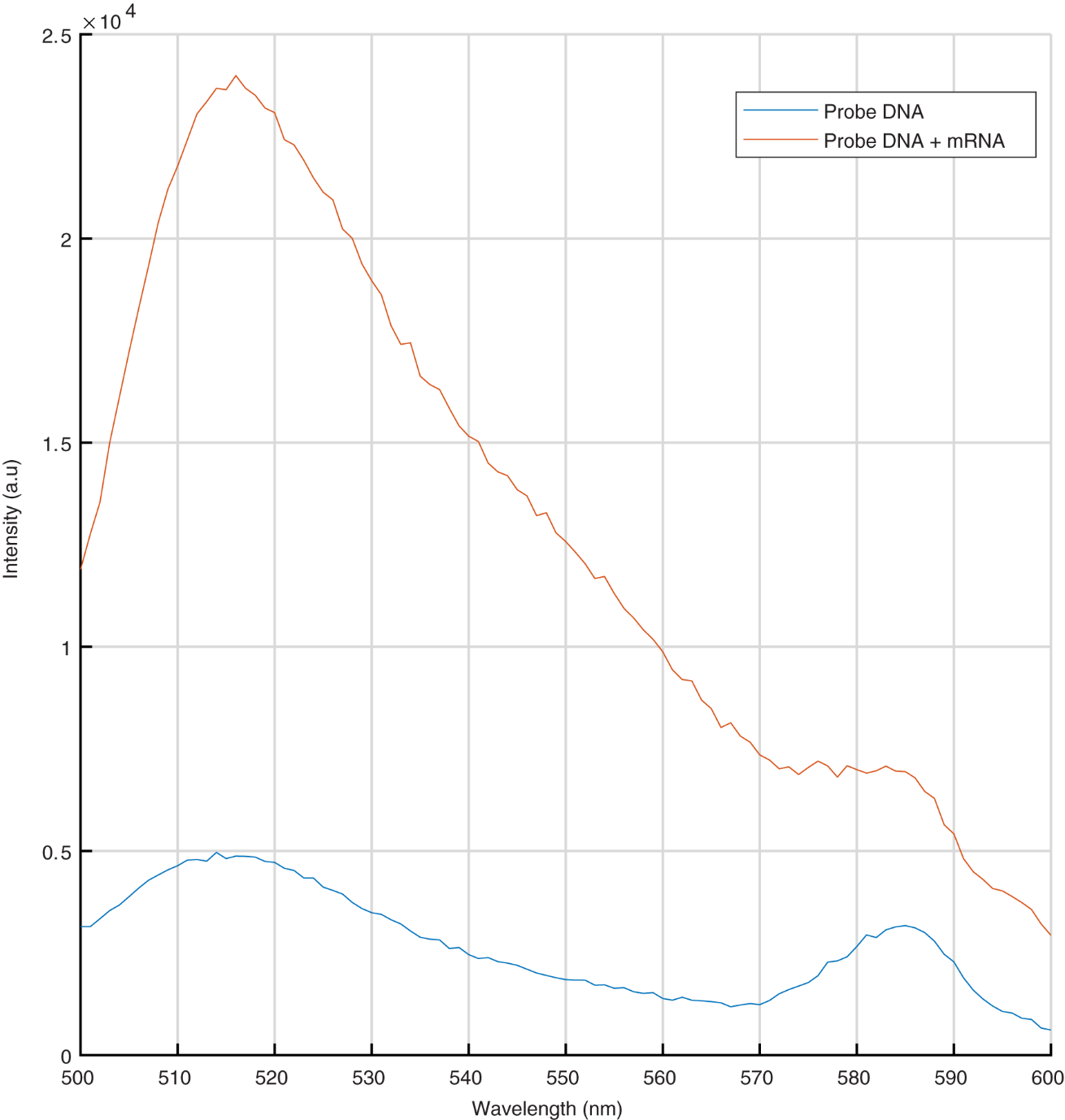

**Supplementary Figure 19.** Ensemble probe DNA binding to mRNA measured with fluorescence spectrometer. Red line: fluorescence of fastFISH probe with mRNA. Blue line: fluorescence of fastFISH probe only.

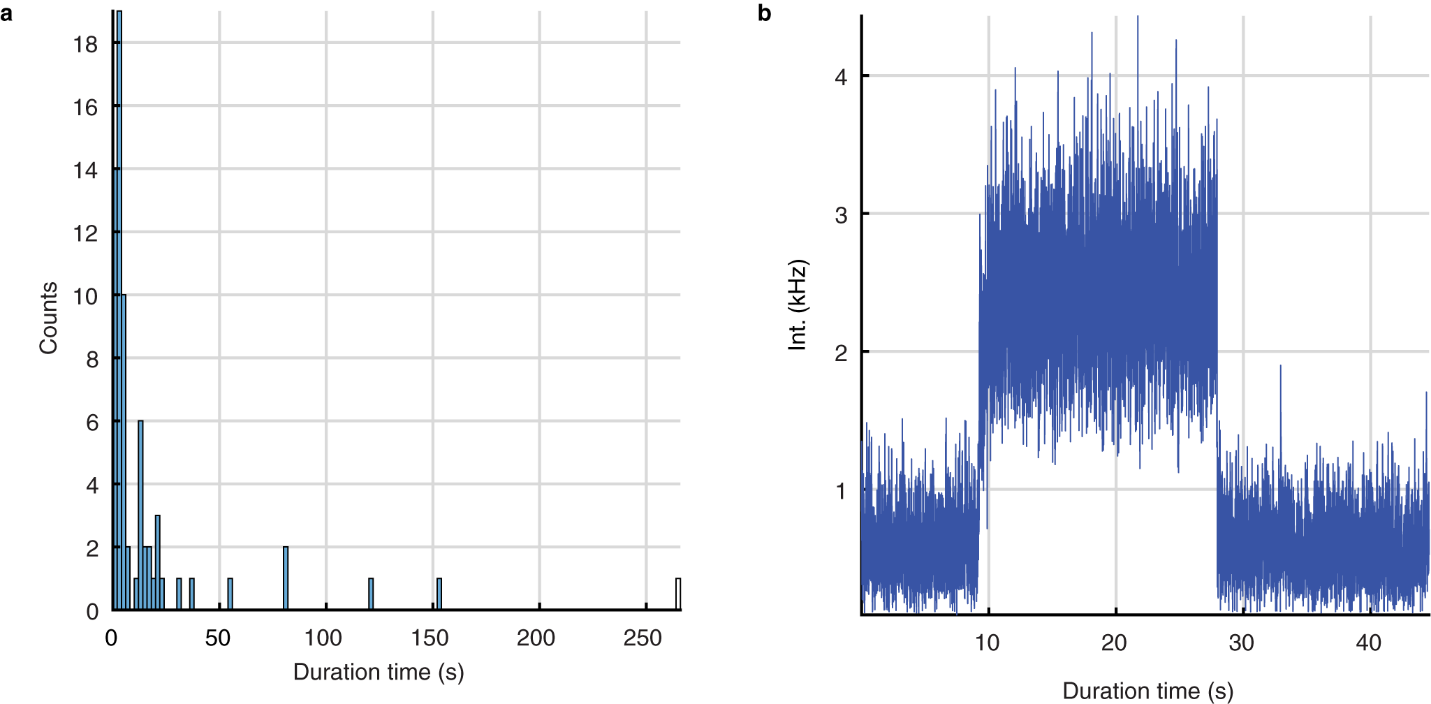

**Supplementary Figure 20.** Alexa 488 bleaching time measurement. (**a**) Histogram of bleaching time of Alexa 488. Bin size: 2s. n=69. (**b**) An example of bleaching events.

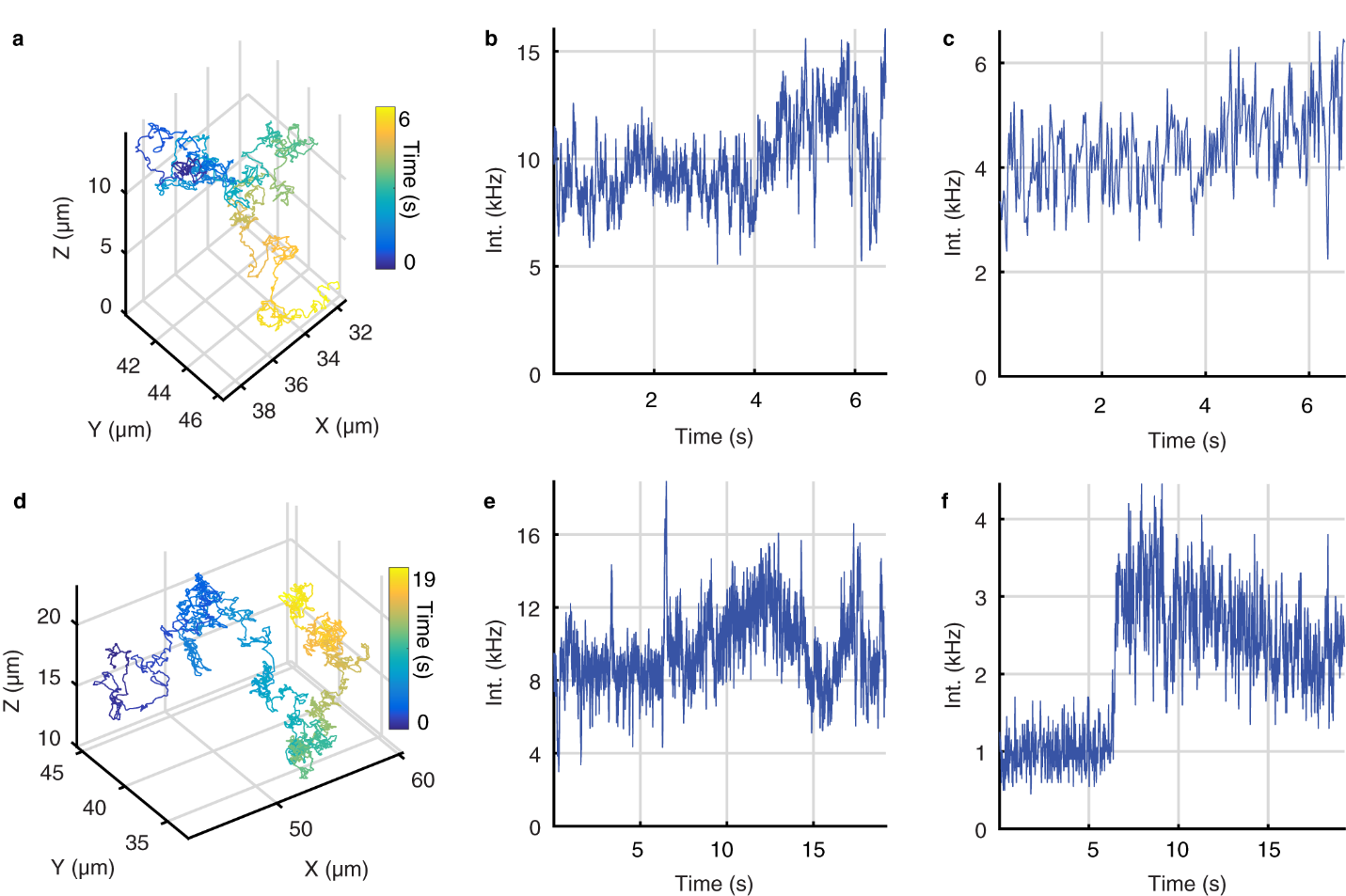

**Supplementary Figure 21.** FastFISH probe DNA non-specific binding to dsDNA in transcription experiment. (**a**) 3D trajectories of Atto 647N labeled dsDNA. (**b**) Fluorescence intensity as a function of time for the trajectory (**a**). (**c**) mRNA channel shows fluorescence signal persisted throughout the trajectory. (**d**) 3D trajectories of Atto 647N labeled dsDNA. (**e**) Fluorescence intensity as a function of time for the trajectory (**d**). (**f**) mRNA channel shows fluorescence increase during tracking that persists to the end of trajectory.

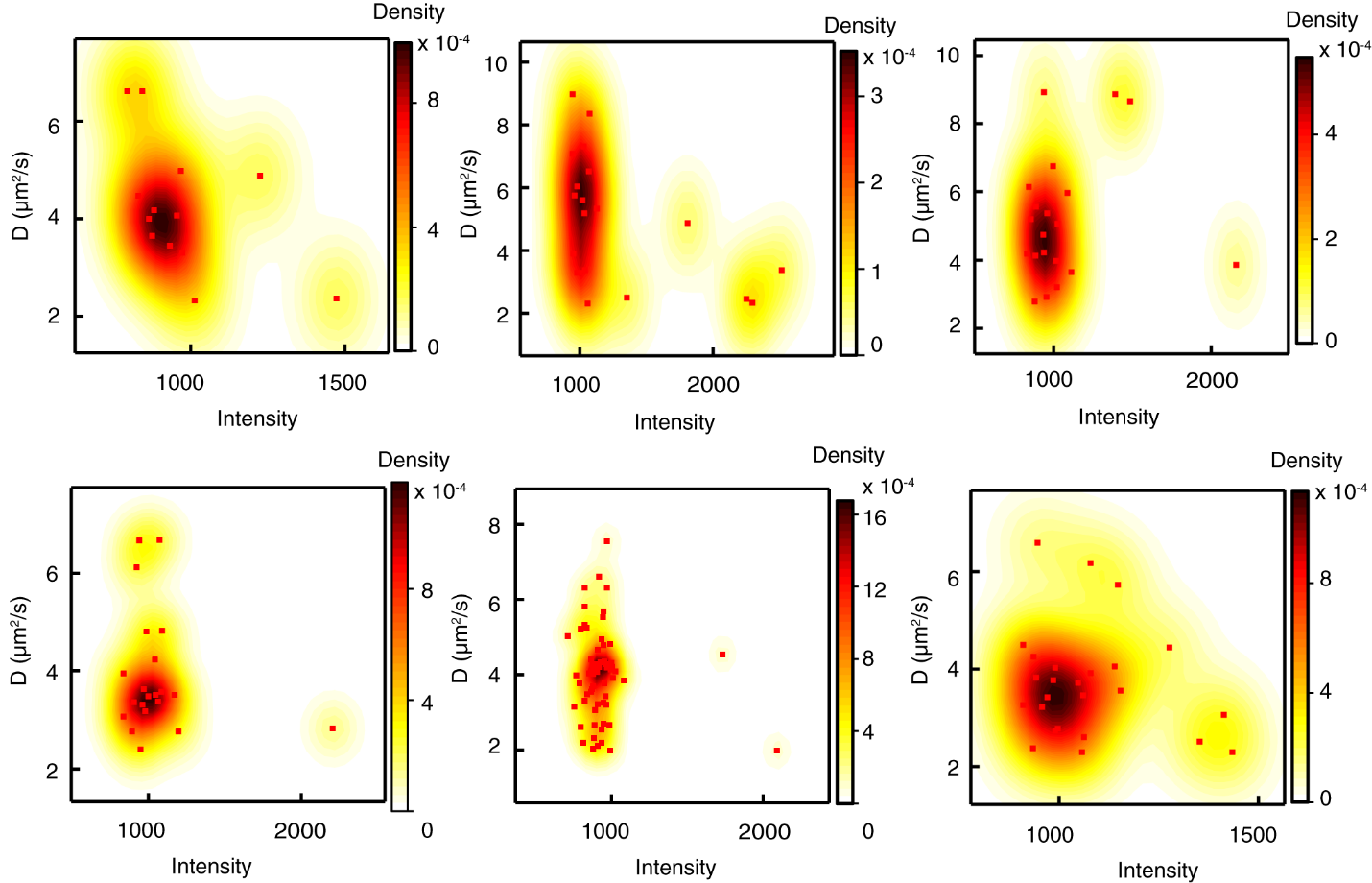

**Supplementary Figure 22.** 2D scatter plot of photon counts of mRNA channel versus diffusion coefficient of template double strands DNA for trajectories in **Figure 4d**. The underlying density map is visualized using 2D kernel density estimation. These scatter plots (from left to right, up to down) are arranged in the same order with **Figure 4d** (from left to right).

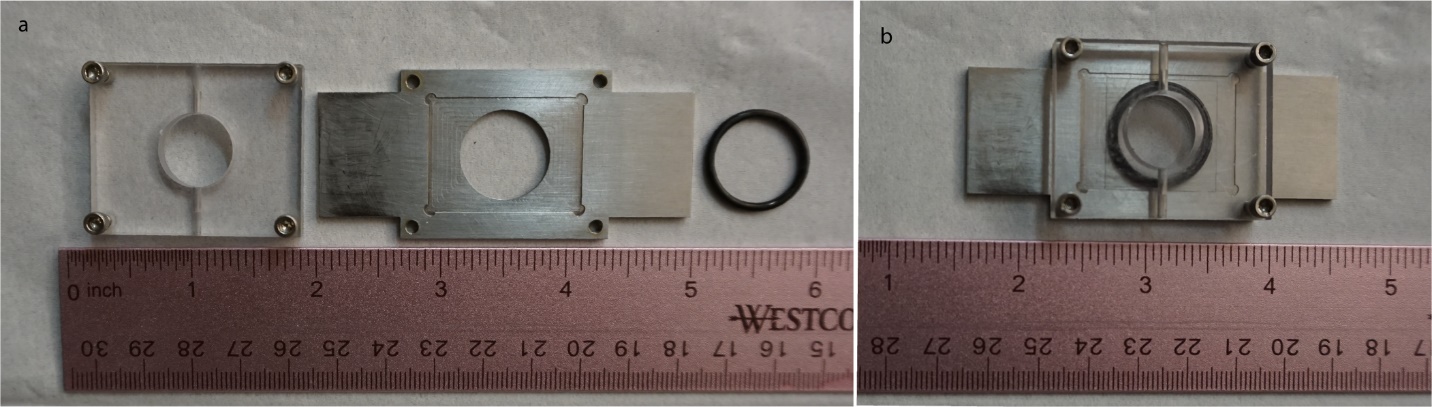

**Supplementary Figure 23.** Sample holder used for 3D-SMART. Sample holder image before (**a**) and after (**b**) being assembled.

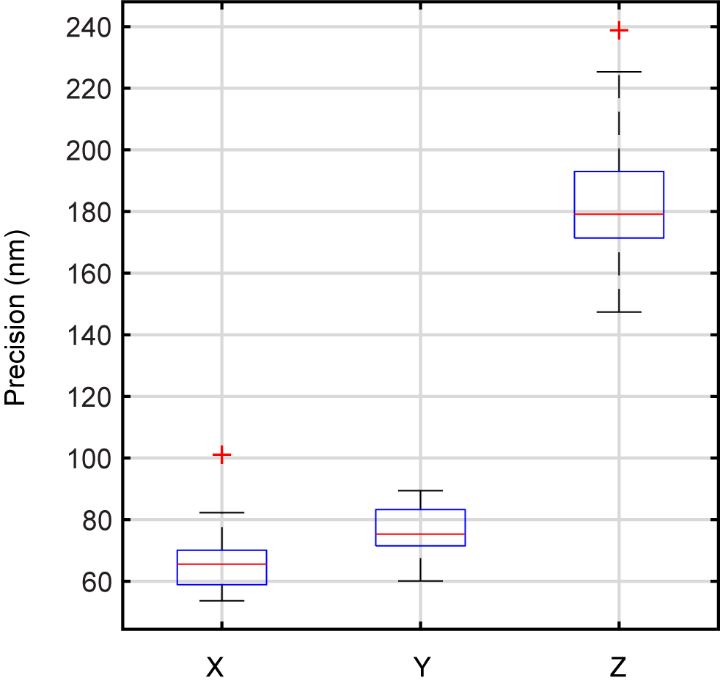

**Supplementary Figure 24.** Tracking precision. The red line represents the median value and the blue box represents the 25th to 75th quartiles of the data. The black whiskers extend to the most extreme data points not considered outliers, and the outliers are plotted individually with red crosses. The mean precision for x, y, and z are 67 ± 12 nm, 75 ± 9 nm, and 184 ± 22 nm, respectively. n = 18.

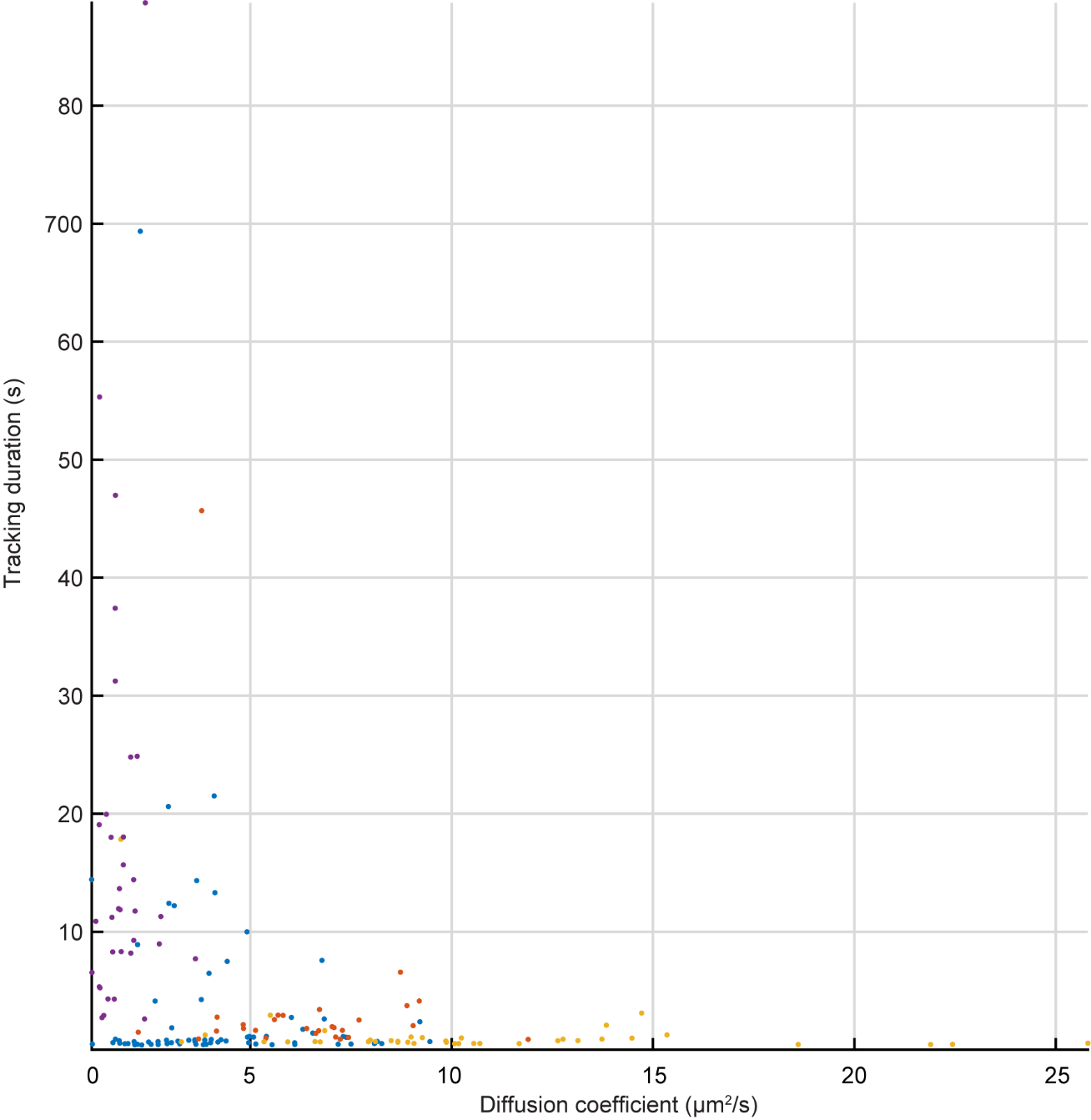

**Supplementary Figure 25.** Scatter plot of diffusion coefficient and tracking duration of dsDNA. Purple dots: 99 bp dsDNA tracked in 70 wt% glycerol. Yellow dots: 99 bp dsDNA tracked in 40 wt% glycerol. Orange dots: 547 bp dsDNA tracked in 25 wt% glycerol. Blue dots: 1385 bp dsDNA tracked in PBS.

| Work | Sample | wt% glycerol | D (μm^2^/s) | Duration (s) |
| --- | --- | --- | --- | --- |
| Wells et al^27^  2010 | Cy5-dUTP | 90 | 3 | 0.100 |
| Han et al^29^  2012 | Rh110 | 96 | 0.43 ± 0.01 | 0.304 |
| Liu et al^31^  2017 | 5’-ATTO633-labeled 8-nt ssDNA | 70 | 4.8 | 0.039 ± 0.036^a^ |
| Keller et al^30^ 2018 | AF488-DNA-AF594 | 90 | 1.24 ± 0.01 | 0.159 ± 0.008 |
| 3D-SMART | ATTO647N | 90 | 2.13 ± 0.55 | 15.997 ± 13.366 |

**Supplementary Table 1**. Comparison of RT-3D-SMT methods on single fluorophores. a - Calculated from the reported geometric distribution fit of the tracking duration with p = 0.13 and a step size of 5 ms.

|  | **Duration (s)** | | | | **Photons (x10^3^)** | | | |
| --- | --- | --- | --- | --- | --- | --- | --- | --- |
|  | **Min** | **Max** | **Mean** | **SD** | **Min** | **Max** | **Mean** | **SD** |
| **Tracking OFF** | 0.265 | 0.856 | 0.457 | 0.169 | 1.783 | 8.475 | 3.408 | 1.494 |
| **Tracking ON** | 0.400 | 52.755 | 15.977 | 13.366 | 3.737 | 634.88 | 193.94 | 161.68 |

**Supplementary Table 2**. Comparison of trajectory duration and photons per molecule with tracking on and off.

**Supplementary note 1：antibunching measurement**

The emission photons were split by a 50:50 beamsplitter and directed to two APDs as showed in Supplementary Fig. 5. The photon arrival time was tagged by TCSPC. The photon pair correlation can be calculated as bellow:

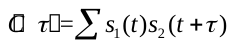

in which s(t) is the detected photon signal at time t for each APD, τ is the lag time. The time resolution of TSCPC is 25 ps. Then the calculated correlation result was binned by 2 ns.

**Supplementary note 2：diffusion coefficient calculation**

The mean square displacement is calculated as below:

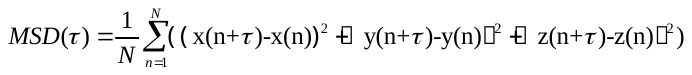

in which x(n), y(n) and z(n) are the coordinates of the trajectory at timepoint n. τ is the lag time. N is the total number of data points that used for calculation. In here we use ¼ of the entire trajectory for MSD calculation. The diffusion coefficient can be obtained by linear fitting of the MSD with lag time τ and the diameter of the particles can be calculated using the Stokes-Einstein relation:

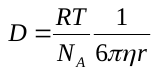
,

in which R is universal gas constant, T is temperature, N_A_ is Avogadro's number, η is viscosity of solution and r is the hydrodynamic radius of particle. For trajectories longer than 20 seconds, only the first 20 s was used for mean-square displacement calculation.

**Supplementary note 3：2D kernel density estimation**

The 2D kernel density estimation map is produced with OriginPro 2017 (OriginLab Corporation). The number of grid points in X/Y for 2D kernel density map in Fig.4e is set to 160 and in Supplementary Fig. 22 is 32. The density at each point (x,y) is estimated by the following equation:

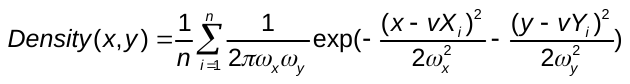
,

in which n is the number of elements in vector vX or vY,
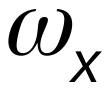
and
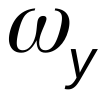
are the bandwidths of X scale and Y scale. Here vX is photon counts of mRNA channel and vY is diffusion coefficient of template double strands DNA.
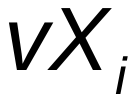
 is ith element in vector vX and
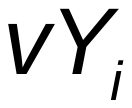
is ith element in vector vY.

**Supplementary note 4：stage calibration**

The maximum voltage readout of the Nano-PDQ piezo stage in the X and Y direction is ~75 µm. However, the stage can still move beyond this range, although the voltage readout is saturated. This saturation was overcome by recording the input command voltage, which was sent to the piezo stage, effectively extending the range of motion to ~95 μm. To do this, we first calibrated the actual stage position with input command voltage by imaging 190 nm fluorescent beads in epifluorescence mode on a sCMOS. The piezo stage was the stepped and the change in the bead position was measured. This calibration was used to recover the particle position when the readout of the piezo stage was saturated (>72 μm).

**Supplementary note 5: Factors that affect collected signal intensity**

There are several factors which will affect the collected signal intensity. The first is the spherical aberration of the high NA objective lens, which leads to decreased collection efficiency as the molecule moves further away from the coverslip. The second is the concentration of glycerol. Glycerol changes the refraction index of the solution, which will affect the fluorescence collection efficiency of the objective lens. Moreover, the viscosity of the solvent can affect the quantum yield of fluorophores through suppression of internal conversion.

Movie descriptions

Supplementary Movie 1.

3D-SMART tracking of a single Atto 647N molecule in 90 wt% glycerol. The top panel shows the molecule’s real-time trajectory and the bottom left panel shows the fluorescence intensity as a function of time. The real-time photon readout at the individual XY laser scan positions is displayed in the bottom right.

Supplementary Movie 2.

Real-time intensity traces with feedback (top) off and (bottom) on. The real-time photon readouts at the individual XY laser scan positions are displayed on the right.

Supplementary Movie 3.

BSA-Atto 647N photo-blinking captured with 3D-SMART tracking. The top panel shows the molecule’s real-time trajectory and the bottom left panel shows the fluorescence intensity as a function of time. The real-time photon readout at the individual XY laser scan positions is displayed in the bottom right.

Supplementary Movie 4.

Real-time tracking of 1385 bp dsDNA tracking in PBS. The top panel shows the molecule’s real-time trajectory and the bottom left panel shows the fluorescence intensity as a function of time. The real-time photon readout at the individual XY laser scan positions is displayed in the bottom right.

Supplementary Movie 5.

DFHBI-1T labeled mRNA tracking in 75 wt% glycerol. The top panel shows the molecule’s real-time trajectory and the bottom left panel shows the fluorescence intensity as a function of time. The real-time photon readout at the individual XY laser scan positions is displayed in the bottom right.

Supplementary Movie 6.

Real-time measurement of transcription on a single, freely-diffusing dsDNA. The top panel shows the dsDNA molecule’s real-time 3D trajectory. The trajectory is labelled with a green sphere when mRNA (fastFISH) signal is observed. The bottom panels show the intensity of Atto 647N labeled dsDNA (red) and the readout of the ssDNA (green). The real-time photon readout at the individual XY laser scan positions is displayed in the bottom right.

**Supplementary codes: Tracking Performance Simulation**

### Simulate Tracking Performance

%This code simulates 2D real-time tracking using experimental piezo response

### Set simulation parameters

tau = 20e-6; %Time step [sec]
N = 1e4; %Number of trajecotry steps
time = tau*(1:N); %create a time vector for plotting

### Laser scan parameters

w = .2e-6; %Beam width
bigW = [w^2 0;0 w^2]; %2D width of the PSF
beamAmp = 0.5e-6; %Laser Scan Size

### Feedback Parameters

K_P = 0; %Proportional Feedback Gain
K_I = 0.012; %Integral Feedback Gain

### Set diffusion and particle parameters

D = 4e-12; %Diffusion coefficient [m^2/sec]
k = sqrt(2*D*tau);

brightness = 6e4; %Particle brightnesss [Hz]
bg = 700; %Background [Hz]

### Generate Trajectory

%Generate steps in x (dx) and y (dy)
dx = k * randn(N,1);
dy = k * randn(N,1);

%Generate XY trajectory
x = cumsum(dx);
y = cumsum(dy);

### Initialize Position Estimation and Stage Variables

%Initial calculated position
x_k_1 = [0;0];

%Initial uncertainty
sigma_k_1 = (0.2e-6)^2;

%Initial stage positions
prevStagePosX = 0;
prevStagePosY = 0;

%Initialize variables
err = [];
stageCommands = [];
x_k = [0 0];
out = [];

### Load impulse response data

measuredImpulseResponse = dlmread('Impulse_Response.txt');

### Track

Loop through trajectory steps

```
for i=1:length(x)

    %Retrieve current beam positions
    [beamX(i),beamY(i)] = kt(i,beamAmp);

    %Correct current laser position with stage
    laserPositionX = beamX(i) + prevStagePosX;
    laserPositionY = beamY(i) + prevStagePosY;
    laserPosition = [laserPositionX laserPositionY]';

    %Get current particle positions
    particlePositionX = x(i);
    particlePositionY = y(i);
    particlePosition = [particlePositionX particlePositionY]';

    %Get number of photons emitted at current step
    emission(i) = poissrnd(tau*emRate(particlePosition-[prevStagePosX prevStagePosY]',laserPosition-[prevStagePosX prevStagePosY]',bg,brightness,bigW));

    %Update step
    x_k = (w^2*x_k_1+emission(i)*sigma_k_1*(laserPosition-[prevStagePosX prevStagePosY]'))/(w^2+emission(i)*sigma_k_1);
    sigma_k = w^2*sigma_k_1/(w^2+emission(i)*sigma_k_1);

    %Error calculation.
    err(i,:) = x_k;
    pkOut(i) = sigma_k;

    %Prediction step
    x_k_1 = x_k;
    sigma_k_1 = sigma_k + 2*D*tau;

    %Store position estimates
    out(i,:) = x_k;

    %Get current stage commands and positions
    stageCom = K_P * err(i,:) + K_I * sum(err,1);
    stageCommands(i,:) = stageCom;
    stagePosX = impulseResponse(stageCommands(:,1),measuredImpulseResponse);
    stagePosY = impulseResponse(stageCommands(:,2),measuredImpulseResponse);

    prevStagePosX = stagePosX(end);
    prevStagePosY = stagePosY(end);
end
```

### Trajectory Statistics

Calculate the total tracking error

trackingError = (sum((x-(stagePosX+out(:,1))).^2)+sum((y-(stagePosY+out(:,2))).^2));

%Calculate emission rate
rate = (sum(emission)/max(time));

%This is the total tracking error per Hz
normalizedTrackingError = (sum((x-stagePosX).^2)+sum((y-stagePosY).^2))/rate;

### Plot Results

figure(1)
subplot(3,1,1)
plot(time,x,time,stagePosX+out(:,1),time,stagePosX)
legend('Particle Position','Calculated Position','Stage Position')
xlabel('Time (s)')
ylabel('X Position (m)')

subplot(3,1,2)
plot(time,y,time,stagePosY+out(:,2),time,stagePosY)
legend('Particle Position','Calculated Position','Stage Position')
xlabel('Time (s)')
ylabel('Y Position (m)')
subplot(3,1,3)
plot(time,emission)
xlabel('Time (s)')
ylabel('Intensity (cts/bin)')

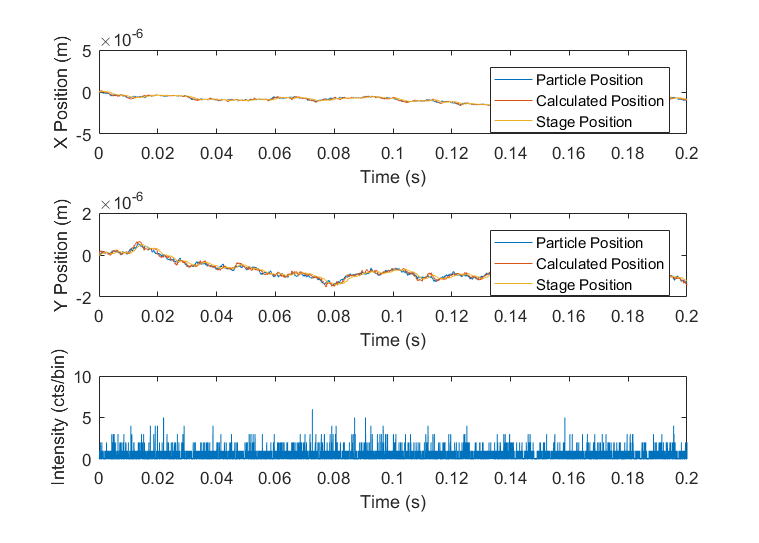

### Impulse Response Function

function out = impulseResponse(x,measuredImpulseResponse)
%Convolve with stage command data
temp = conv(x,measuredImpulseResponse);
%Crop data
out = temp(1:length(x));
end

### Knight's Tour Coordinates

function [xOut,yOut]=kt(index,d)
x = d*[-1;-0.5;-1;0;1;0.5;1;0;0.5;1;0;-1;-0.5;0.5;1;0.5;-0.5;-1;0;1;0.5;-0.5;0;-0.5;-1];
y = d*[1;0;-1;-0.5;-1;0;1;0.5;-0.5;0.5;1;0.5;-0.5;-1;0;1;0.5;-0.5;-1;-0.5;0.5;1;0;-1;0];
xOut = x(mod(index-1,length(x))+1);
yOut = y(mod(index-1,length(x))+1);
end

### Emission

function out=emRate(x,c,b,s,w)
%x - position of particle
%c - position of beam
%b - background
%w - spatial covariance
out=s*exp(-0.5*(x-c)'*inv(w)*(x-c))+b;
end

[*Published with MATLAB® R2016b*](http://www.mathworks.com/products/matlab)

Supplementary Data (separate files)

Impulse_response.txt: Impulse response data from piezoelectric stage used to model tracking performance.

model1track.m: MATLAB tracking simulation code.
